# DOCUMENTING AND EVALUATING DATA SCIENCE CONTRIBUTIONS IN ACADEMIC PROMOTION IN DEPARTMENTS OF STATISTICS AND BIOSTATISTICS

**DOI:** 10.1101/103093

**Authors:** Lance A. Waller

## Abstract

The dynamic intersection of the emerging field of Data Science with the established academic communities of Statistics and Biostatistics continues to generate lively debate, often with the two fields playing the role of an upstart (but brilliant), tech-savvy prodigy and an established (but brilliant), curmudgeonly expert, respectively. Like any new discipline, Data Science brings new perspectives and new tools to address new questions requiring new perspectives on traditionally established concepts. In this paper, we explore a specific component of this discussion, namely the documentation and evaluation of Data Science-related research, teaching, and service contributions for faculty members seeking promotion and tenure within traditional departments of statistics and Biostatistics.

## INTRODUCTION

The development of the field of Data Science has generated much discussion, enthusiasm, and investment within Colleges and Universities in recent years. Within the general field of Statistics, Data Science has extended and expanded concepts such as significance testing and survey sampling to address inference in unstructured, big data settings, but, at the same time, some high-profile overviews of Data Science fail to mention the field of Statistics at all, despite its fundamental role in the process (see, e.g., Davidian and Louis, 2012). While recent reports have examined the need for statistical thinking within Data Science research and training (National Research Council 2013, 2014), fewer have addressed the value of Data Science concepts within the field of Statistics, particularly with respect to the recognition of Data Science research, teaching, and service activities by faculty members in traditional Statistics and Biostatistics academic departments.

As in the development of and definition of any new area of scientific inquiry, leading researchers and instructors in the area of Data Science typically are not themselves trained as data scientists, rather they brought their own training and experience from computer science, mathematics, statistics, and other fields to the table to help define the scholarly elements of research, teaching, and service for an emerging science. These definitions remain fluid but play a critical role in determining evidence of academic success in each arena.

Several points of view are relevant in this discussion. From the perspective of a junior faculty member in the field, it can be invigorating to be involved in the growth of a new field, but also concerning to wonder if contributions to interdisciplinary science will be appropriately valued and appreciated as scholarly contributions when one is evaluated for promotion and/or tenure within traditional disciplinary departments. From the perspective of a Department Chair seeking to support meaningful participation in the development of emerging areas of science clearly relating to Statistics and Biostatistics, it can be exciting to encourage leadership in cutting edge development of a new field of inquiry, but it can be challenging to determine how best to package such accomplishments to ensure full appreciation of an individual junior faculty member’s unique accomplishments within each step of the review process.

The sections below seek to address these related sets of concerns in the particular setting of evaluating Data Science contributions for promotion and tenure review within research, teaching, and service in traditional departments of Statistics and Biostatistics. We begin with a brief and generic overview of the academic promotion process including a review of the typical dossier that serves as evidence for review at each step of the process. We then outline the typical components of the dossier with suggestions as to where and how to incorporate, document, and highlight contributions to Data Science, noting the specific need to establish interpretable context for these contributions. Like the field of Data Science itself, some of these elements are dynamic and are likely to change (perhaps rapidly) over time so we hope this overview encourages ongoing discussion on the topic among junior and senior faculty members, department chairs, and academic administrators.

## THE ACADEMIC PROMOTION PROCESS

Roughly speaking, the academic promotion process resembles a pre-Copernican view of the universe centered on the junior faculty member, expressed as a set of concentric administrative layers at the Department, School/College, and University level. The promotion review process passes outward from the individual through each of these layers. Generally, the process is similar for both tenure-track and non-tenure-track faculty members where the tenure track often involves stricter time constraints, a clear up-or-out decision, and may sometimes involve slightly more comprehensive documentation of accomplishments.

The first step for any promotion begins early, often during the candidate’s initial interview for an academic position. It is always beneficial for the candidate to begin an ongoing conversation regarding the process, expectations, and timeline associated with promotion. This begins an essential and ongoing conversation between a Department Chair and a faculty member, but it is also wise for a candidate to discuss the process with other senior faculty within the department who will be evaluating their accomplishments, departmental representatives on the School or College Promotion and Tenure committee, as well as recently promoted faculty members to gain perspective regarding their experiences with the process. These conversations will introduce the candidate to the process early and often will provide a mechanism for initiating dossier documentation very early in the candidate’s career allowing regular updates rather than a flurry of preparation immediately before a promotion review.

The formal promotion evaluation process begins focused on the individual junior faculty member seeking promotion through her/his accomplishments in and contributions to the candidate’s area of expertise, to the candidate’s Department, to the candidate’s School, to the candidate’s University, and to the candidate’s profession. We typically summarize these accomplishments as contributions in each of three areas: Research, Teaching, and Service. My own institution requires individuals be evaluated as “Excellent” on one of these three areas (with specified criteria for excellence provided in the guidelines for promotion), and evaluated at least as “Very Good” in the other two.

In order to initiate the promotion process, the candidate works with the Department Chair to prepare a dossier for review at the following stages, typically in the following order:

1. An informal review by Department faculty currently at or above the level to which the candidate seeks promotion (e.g., tenured faculty in the case of an individual being evaluated for tenure, Professors for those seeking promotion to the rank of Professor). These faculty review a full curriculum vita (CV) and 2-3 page Personal Statements (see below) by the candidate regarding their past performance, current efforts, and future plans in each of the Research, Teaching, and Service areas.
2. A set of several (typically six) individual evaluations, summarized in formal letters, by external experts working in areas similar to the candidate’s areas of expertise but not working closely with the candidate (i.e., typically not publishing or directly collaborating with the candidate). Depending on the status of open records laws and policies associated with the home institution, these letters may or may not be held as confidential.
3. A formal review and vote by the faculty members in step 1, based on the updated dossier material (the CV, the Personal Statements, and the external letters from step 2). The Department Chair will conduct the vote and summarize results to pass along to the next step along with the dossier materials.
4. A formal review and vote of approval by a Promotions and Tenure committee within the academic unit overseeing the Department (often a School or a College);
5. Review and approval of the Promotions and Tenure committee’s recommendation by the Dean of the School or College;
6. Many Universities include a review by an University-level advisory committee reporting to the President; and
7. Final review and approval (or not) by the University Board of Trustees (or similar governing body).

At each stage of the review process, the results and documentation of each previous step are summarized and included in evaluation materials, e.g., the results of votes at the Department and School level are summarized in a letter by the Department Chair and included for review at subsequent levels of the process.

In addition to the above promotion review, many universities also conduct an interim review of the progress of junior faculty, e.g., during year 3 of the six year probationary period for tenure-track Assistant Professors, or annually for some universities). Such interim reviews incorporate elements of steps 1-4 in the outline above (typically with the exception of the external review by experts). These interim reviews provide an excellent opportunity for junior faculty to keep their dossier items up to date and, importantly, to gauge the reception of accomplishments and plans for future directions by those faculty members who will be voting on promotion within the candidate’s Department and School. The interim reviews also provide an opportunity to assess whether the candidates accomplishments are being communicated in a manner that is fully understood and appreciated by reviewers at these critical early steps of the process.

It is important to note that evaluations at the first few stages in the process above typically focus on the individual’s contributions within a disciplinary environment (i.e. addressing the question “is the candidate a good academic statistician?”), and later stages by reviewers further removed from the individual’s area of expertise who rely more on general assessments of academic accomplishment and summaries from earlier stages of review (i.e., addressing “is the candidate a good faculty member?”). The interdisciplinary nature of Data Science requires a candidate to prepare carefully for this typical (but often underappreciated) feature of the review process, i.e., by preparing strong, documented evidence to provide positive responses to both questions. This is especially important if many of the candidate’s accomplishments fall outside of the “typical” Research, Teaching, and/or Service familiar to the senior faculty in the candidate’s Department or to senior faculty in the candidate’s School or University. It is also particularly important for the candidate to have conversations with the Department Chair, senior faculty in the Department, and the Department representatives to the Promotion and Tenure Committee in order to assess (a) their receptiveness and appreciation of interdisciplinary efforts in assessing promotion, and (b) what sorts of evidence they find most convincing in evaluating excellence in such contributions.

## DIFFERENT TYPES OF SCHOLARSHIP

In the 1990s, Earnest Boyer of the Carnegie Foundation for the Advancement of Teaching provided an important resource promoting the value of different types of interdisciplinary scholarship through his publication of *Scholarship Reconsidered: Priorities for the Professoriate* (latest, expanded edition: Boyer et al. 2016). This report, highly referenced by university administrators such as deans, provosts, and presidents, but often less well known by junior faculty, outlines the value of multiple types of research within academia, specifically noting four types of scholarship. The first, *scholarship of discovery*, mirrors the standard disciplinary model of original research advancing knowledge within a field, often evidenced by peer reviewed publications in established disciplinary journals and success in obtaining competitive research funding. A second type, *scholarship of integration*, recognizes the synthesis of information across traditional disciplines, across subdivisions within a discipline, or across time. Such scholarship creates new knowledge by bringing novel links between specific concepts, tools, and studies from disparate fields of inquiry. Boyer’s third type, *scholarship of application* (sometimes called *scholarship of engagement*), goes beyond simply applying existing tools (as would a technician) to value the deep collaborative contributions of experts in disciplinary knowledge in creating advances in interdisciplinary studies, particularly within a team science framework. The fourth type, *scholarship of teaching and learning*, values the systematic study of pedagogical methods for the transfer and creation of new knowledge between faculty members, colleagues, and the next generation of scholars.

These four categories provide rich support for many current efforts within the field of Data Science and its link to academic departments of Statistics and Biostatistics. Clearly the scholarship of discovery recognizes disciplinary advances and is documented by traditional peer-review publishing and competitive grant funding. The scholarship of integration is immediately extensible to Data Science, particularly with respect to linking new types of data and developing new analytic tools, hence enabling new lines of inquiry. The scholarship of application is evidenced by interdisciplinary publishing, the creation of data repositories and complex data sets, and clear contributions unique to the candidate within an interdisciplinary team. Finally, the rapid development of training programs, concentrations, and degree programs within the area of Data Science offers multiple opportunities for the scholarship of teaching and learning. We should note, however, that success in this area extends well beyond simply teaching new courses and advising students, it involves research and discovery on the modes and methods of instruction and learning, an area of clear interest in the statistical education research community, but only just developing in the broader area of Data Science.

## DOCUMENTING ACADEMIC ACCOMPLISHMENTS AND PROVIDING CONTEXT FOR DATA SCIENCE CONTRIBUTIONS

We next examine in detail the core set of evidence to be evaluated for promotion, i.e., the elements of the individual’s promotion dossier, a packet of information typically including (at least) three components: (1) a Personal Statement (or Statements) by the candidate summarizing their contributions to and future plans in the areas of Research, Teaching, and Service; (2) a full curriculum vitae (CV) summarizing the individual’s accomplishments to date; and (3) a set of external evaluation letters. The dossier often also includes a set of a handful of representative publications illustrating contributions to Research, teaching evaluations and sample syllabi illustrating contributions to Teaching, and a full summary of contributions to Service. Different institutions offer slight variations to this general framework, but the elements listed above are fairly consistent across most universities in the United States.

Applying these general guidelines to the specific area of Data Science, we next review each component of the dossier in detail.

### Documentation in the Personal Statement(s)

In the Personal Statement(s) (either a single statement covering Research, Teaching, and Service or three separate statements), the candidate provides a summary of academic accomplishments in each area, but the candidate also places these accomplishments in context of her/his broader professional progress. That is, the Personal Statement offers the candidate not only a forum to highlight and summarize accomplishments but to also to express why these contributions matter and where these contributions are leading the candidate and the field of interest more broadly. As noted above, it is important for the candidate to provide evidence of past success and outline a trajectory for future directions. A strong Personal Statement will illustrate how the candidate has built on past work in the field, illustrate the candidate’s participation in present initiatives (their own and more generally), and illustrate the candidate’s vision of future directions for themselves and their area of inquiry. The Personal Statement also offers a place to define the candidate’s contributions within the framework of Boyer’s different types of scholarship, particularly with respect to the scholarship of integration and the scholarship of application. Finally, the Personal Statement offers a the candidate the opportunity to discuss accomplishments from two important perspectives: the candidate’s *contributions*, i.e., how the candidate’s work contributes to the field; and (equally important, but often overlooked) the *candidate’s* contributions, i.e., how the work is dependent on the candidate’s unique involvement, not just the involvement of anyone in the field. This second perspective is particularly relevant for interdisciplinary contributions within Data Science. Framing accomplishments in this way offers the candidate the opportunity to highlight their personal contributions to projects shared within broader research teams in order to make the case that the accomplishments required the participation of this particular candidate and their skills, not simply the presence of anyone with statistical skills.

### Data Science Context in the Personal Statement(s)

The Personal Statements offer the candidate the opportunity to place their research, teaching, and service into the broader perspective of her/his professional vision: the motivation for their work, the types of contributions she/he makes individually and as part of a collaborative team, and the types of things the candidate hopes to do in the future. This is an excellent opportunity for the candidate to define a personal view of what constitutes Data Science, what requires innovation, what the candidate has done in this regard to date, and what opportunities present themselves for future work. The Personal Statements are the perfect place for the candidate to provide a personal definition of Data Science, to identify how Data Science links to but differs from many traditional paths in academic Statistics and Biostatistics, and to articulate why this matters. A clear general definition with specific examples mentioned in each of the Research, Teaching, and Service sections will frame the discussion for reviewers at all stages of the process, since the external reviewers, voting faculty members, and upper level administrators will all have their own personal perspectives on emerging areas of inquiry at varying levels of specificity. The Personal Statements are an opportunity for the candidate to define the discussion rather than hope their accomplishments will be clearly evident at all levels of review.

### Documentation in the CV

The next element of the promotion dossier is the candidate’s full curriculum vita (CV), typically providing background education, employment history, awards and honors, as well as a full list of peer-reviewed publications and grants, conference and seminar presentations, teaching/advising/mentoring activities, and service to the Department, School/College, University, and profession. In the CV, reviewers seek evidence of research success through peer-reviewed publications, competitive grant funding, and invitations to speak at conferences and seminars. For teaching success, reviewers typically look for good and continually improving teaching evaluations, growing course responsibilities, innovations and new ideas in the classroom, and self-awareness evidenced through the development and articulation of an overall teaching philosophy by the candidate. For service, reviewers look for participation on committees at the Department, School/College, and University level as well as participation in activities associated with various professional organizations, refereeing and membership on editorial boards, and participation in research and grant review panels (e.g., for the National Science Foundation or the National Institutes of Health, etc.).

### Data Science Context in the CV

The traditional academic CV in the fields of Statistics and Biostatistics typically highlights research contributions through peer-reviewed publications, especially those in journals with strong reputations in the field, and grant funding, often as Principal Investigator. Many times, research contributions in Data Science result in the creation of novel software tools, merged and curated data sets, and contributions to research teams where the candidate plays an essential role, but may not serve as the lead investigator. It is critical for the candidate to work with her/his Department Chair to provide context for these contributions and to raise awareness of the value of such contributions among the senior faculty in the Department, on the Promotion and Tenure Committee, and at higher levels of the promotion process. Some of this can be accomplished through ongoing conversations, and, as noted above, the candidate can provide important context in the Personal Statements, but it is equally important to document the accomplishments within the CV to make sure they are noticed and appreciated.

As a faculty member in a department of Statistics/Biostatistics, there will be an expectation of some publications in traditional journals in these fields, but the candidate will likely also have publications in the area of Data Science. Publications particular to Data Science likely will appear in newer, electronic journals rather than long-standing established journals. The candidate should note the relevance and reputation of the journals, and include new metrics of impact for her/his publications such as “most downloaded” or “highly cited” over a period of time, as provided by some newer, online journals. The candidate should be careful to distinguish peer-reviewed publications from non-peer reviewed publications. It is fine to list non-peer-reviewed publications, but these should be listed in a separate, clearly marked section. In addition, some computational fields place more stress on refereed meeting proceedings than on traditional journal publications, due to timeliness and competitiveness of review. If the candidate has publications appearing in such proceedings, a parenthetical note identifying the acceptance rate can be helpful for reviewers who may otherwise view proceedings as a non-peer-reviewed publication.

Software development and data wrangling will likely be a key aspect of any candidate involved in Data Science, and typical CVs in Statistics/Biostatistics do not consistently highlight such contributions. One common approach is to have a separate section for software tools and possibly for curated data, but the notion of peer-review and citations for software and data are not clear, nor are they consistently used but new options are rapidly becoming available. Software (e.g., R packages) may be accompanied by peer-reviewed publications in journals such as *Journal of Statistical Software*, but may also be closely tied to peer-reviewed journal or proceedings articles already appearing in the candidate’s publication list (e.g., an R package associated with the statistical methodology proposed in a publication in a statistical journal). If the later is the case, I suggest a parenthetical comment linking the software package to the associated publication (or publications). This allows the candidate (and the Department Chair) to highlight citations as well as downloads and provides a link between the software tool and the peer-reviewed publication that may be helpful for more traditional reviewers.

While some areas of science have well documented data repositories and mandates exist for making federally funded research project data publically available, two challenges remain in the general recognition of data, particularly complex, linked, and curated data, as citable, scholarly research output. The first is the establishment of a peer-review equivalent to publications for quality control, and the second is the absence of a standard method to cite complex data sets (including verifiable attribution and date/version labels). Some citation standards certainly exist, but, in general, these are not (yet) universally applied and the true impact of a data set likely falls somewhere between the full number of downloads and the current number of formal citations. These two issues present an obstacle for the clear recognition by reviewers (both internal and external) of the impact of the contributions of data wrangling and curation to the advancement of Data Science as well as Statistics/Biostatistics.

Some recent developments in this regard include the work of the Research Data Alliance (and their Data Citation Working Group, https://rd-alliance.org/groups/data-citation-wg.html, providing guidelines for data citation of evolving data sets), among other groups, and the establishment of the Joint Declaration of Data Citation Principles (Data Citation Sythesis Group, 2015, https://www.force11.org/group/joint-declaration-data-citation-principles-final). Both represent recent developments in establishing guidelines for data citation. The Data Citation Principles include: importance (i.e., “data should be considered legitimate, citable products of research”), credit and attribution (e.g. are all individuals involved credited with the data?), evidence (i.e., when results rely on specific data sets, these data should be cited), unique identification (which version of the data are you using?), access (i.e., are the data available?), persistence (i.e., past data should be available as well as updates), specificity and verifiability (e.g., does the citation clearly identify which version of a data set was used?), and interoperability and flexibility (e.g., data access should work across platforms). These are important steps, but equally important to their definition will be the adoption of the principles in practice by the scientific community, including the fields of Statistics and Biostatistics. As a side note, it is and will continue to be important for statisticians to be involved in these activities and continued developments in developing data citation guidelines and practice.

Currently, many details regarding the process of creating and curating complex data sets are deferred to the supplemental information section of a publication, often receiving much lighter review than the main body of the paper but containing information critical for the reproducibility of the results. In response, some novel publication outlets are appearing which provide researchers the opportunity to submit their data *and* details on its construction for peer review and publication, in parallel with the original research report. These opportunities provide a unique, citable, digital object identifier (DOI) for (i) the original paper, (ii) the detailed description of data development, and (iii) the data themselves. Two examples of this approach include the Dryad Digital Data repository and an online journal by the Nature Publishing Group, *Scientific Data*. These are simply two examples and similar outlets are also available and in development. The Dryad Digital Data repository (http://datadryad.org/) provides a digital repository for data that meet most (if not all) of the Data Citation Principles listed above, again providing a DOI associated with the data and a link to the original publication. *Scientific Data* (http://www.nature.com/sdata/) takes things a step further and publishes peer-reviewed “data descriptors”, full-length papers about the data development process spelling out the details that are often relegated to supplemental information but are essential for documenting the elements of Data Science involved in the creation of the data set, again published with a DOI. These two examples and others like them, provide an outlet for data-oriented research output that are more similar to traditional publishing measures (peer-reviewed and citable) and more familiar to reviewers from traditional disciplinary areas.

As a final consideration, newer sources of research writing such as social media posts and blogs do not fit neatly into the traditional promotion dossier, but can have a documented impact on the field. Such activities should be noted in the CV in a clearly identified section along with documentation of impact. Again, the Personal Statement allows the candidate to provide context for this information and parenthetical notes may clear up potential misconceptions. The key element will be to provide evidence of the influence by the candidate on the field through such activities. The candidate and Department Chair should discuss ways to document and demonstrate such impact through reports of reposts or through identification of external reviewers who can be asked to specifically comment on these elements in their evaluation letters.

The CV also provides evidence of accomplishments in Teaching. As noted above, such evidence includes standard documentation (lists of courses developed and taught, lists of enrollment, sample syllabi, and summarized teaching evaluations). Reviewers typically examine this information for evidence of dedicated effort associated with training the next generation of scholars within the candidate’s own discipline, but also in training future collaborators the key elements associated with statistical thinking. The emerging nature of Data Science provides opportunities for generating new courses or revised curricula for existing courses to weave elements of computation, data management, markup reports, and data wrangling into courses at all levels based on research trends and demands of future employers (National Research Council 2014). As with Research, the Personal Statement provides the candidate the opportunity to place the full Teaching portfolio in the broader context of the candidate’s professional goals and accomplishments. The emphasis on computation and technology within Data Science also opens the door for online instruction ranging from YouTube channels devoted to instruction in specific software packages, to massive, open, online courses (MOOCs) enrolling thousands of students. These new opportunities for instruction may be unfamiliar to traditional reviewers (internal or external), and may require additional context and metrics to document impact and influence on the field. For example, MOOCs are notorious for having very large initial enrollments with a low completion rate, and it is important to be up-front about the full picture of such activities to stem potential skepticism by reviewers. Linking YouTube instruction to citable summaries, or particular software packages or publications can also help. As with novel sources of publication (e.g., blogs) mentioned above, the candidate should work closely with the Department Chair to document and demonstrate impact on the field and identify potential reviewers who can speak direction to this impact.

Finally, the CV should highlight Service contributions including traditional lists of committee membership, refereeing activities, editorial board service, and participation in meeting organization and professional organizations. Leadership roles can follow committee participation and both levels of activity are important to highlight. Data Science related activities in this standard list should be identified in the Personal Statement (e.g., serving on an organizing committee for a Data Science workshop or organizing invited speaker sessions on Data Science topics at both Data Science and statistical conferences). Candidates with Data Science activities may demonstrate refereeing or associate editor duties for journals in both Data Science and Statistics/Biostatistics or referee activities for data publishing outlets (e.g., those mentioned above). Many Schools, Colleges, and Universities are in the midst of strategic planning activities relating to Data Science, Big Data, etc., and Department Chairs are often looking for representatives with knowledge in both Statistics/Biostatistics and Data Science to serve on such committees. These committees also provide valuable networking and collaborative opportunities for junior faculty, and often provide leadership opportunities down the road. Candidates and Department Chairs alike should be vigilant for such opportunities but also careful to weigh the potential benefits to the candidate against the associated time, effort, and likely outcomes involved. These activities may be aggregated as “Data Science Service” in the CV or simply interspersed among other Service contributions, but, again, should be highlighted as targeted activities in the Personal Statement.

### Documentation in the External Letters

The external letters are an essential element of promotion review and consist of three separate components: the letter writer, the letter content, and the letterhead. We consider each of these in turn.

The *letter writer* should be an established and successful academic who can write knowledgeably about the candidate’s accomplishments and evaluate overall success in Research, Teaching, and Service. The letter writers are typically “arm’s length” evaluators and have had limited direct collaboration with the individual. It is sometimes a challenge for Department Chairs to identify individuals who know the candidate’s work well but have not (yet) collaborated with the candidate. Candidates can provide some suggestions of individuals they feel well suited to evaluate accomplishments, similarly, candidates can request that particular individuals with perceived conflicts-of-interest not serve as external reviewers. That said, most, if not all, institutions require some external reviewers identified independently of the candidate’s suggestions. Therefore, a candidate should provide a representative but not comprehensive list of potential knowledgeable external reviewers. In addition, for candidates with a Data Science focus, it is important to clearly communicate to the Department Chair the *type* of reviewer (e.g., individuals who make use of large-scale distributed computing) who would best be able to appreciate and evaluate contributions to Data Science, to aid in identifying additional reviewers.

The *letter content* provides detailed assessment of the candidate’s accomplishments and career trajectory. Some universities require explicit comparison to others in the candidate’s general stage of career as well as an evaluation as to whether the candidate would be competitive for promotion at the letter writer’s home institution. Other universities prohibit such statements and it is helpful for candidates to know the rules of their own institution. Letter writers are often experts in areas related to the candidate’s work and can provide disciplinary details relating to the quality of journals, competitiveness of grants, success in instruction and mentoring, and service activities, as well as broader contributions to the candidate’s Department, School/College, University, and profession.

Including the *letterhead* as component of the evaluation letter may seem a bit cynical or facetious, but the reputation of both the letter writer and the letter writer’s home institution both carry weight in the evaluation, particularly as the dossier passes through the latter stages of review. As the candidate’s promotion advances through the levels outlined above, the individuals reviewing the materials will be less familiar with the candidate’s particular field of study, target journals, and funding agencies and will rely more on broader measures of assessment and give weight to qualifications of the external reviewers such as titles (e.g., Distinguished Professor, Department Chair) and the reputation of their home institution (e.g., is this a “peer institution” the candidate’s institution considers an upstart, an equal, or an aspiration?). It is very helpful for letter writers to come from institutions that have demonstrated success in the areas relating to the candidate’s accomplishments and goals.

### Data Science Context in the External Letters

External reviewers are encouraged to provide their view of the candidate based on the dossier as well as any personal experience they have interacting with the candidate’s work. The context provided by the Personal Statements and the organization of the CV will be critical in presenting the contributions of the candidate in light of their impact on the candidate’s discipline (Statistics/Biostatistics) as well as on Data Science. The organization of material can be extremely helpful in documenting the vision, experience, and trajectory of the candidate in each of Research, Teaching, and Service. The candidate has the opportunity to frame material to illustrate connections and the overall career path, and the Department Chair has the opportunity to request specific comments relating to the integration of novel Data Science elements within traditional disciplinary accomplishments. It is not difficulty to identify candidate’s strengths within a well organized dossier, but it can be a challenge if the relevant items are not linked together.

## SUMMARY

In summary, it is in the best interest of both the candidate and the Department Chair to put together an accurate and organized assessment of the candidate’s accomplishments in Research, Teaching, and Service within a dossier where the elements intertwine to make the strongest case possible for promotion. The Personal Statement(s) provide personal context of past work, vision, and future plans, allowing the candidate to place Data Science accomplishments in perspective within their professional development in a traditional disciplinary environment. The CV documents accomplishments in both traditional areas (e.g., publications, grant support, classroom teaching, committees), as well as emerging outlets for scholarly productivity (e.g., DOIs for data, data descriptions, blogs, online education). The external letters provide the perspective of the broader academic community and reflect the letter writer’s perception of the candidate’s accomplishments within current and future trends of academic success. Together, the elements of the dossier provide the basis for review of the candidate’s progress by senior faculty and administrators in the candidate’s Department, School/College, and University.

While promotion considerations should not be an obsession for junior faculty, I do feel it is very helpful to be familiar with the local steps of the process and the local rules governing promotion at the candidate’s institution. I suggest early and frequent discussions between the candidate and Department Chair (e.g., during annual reviews) beginning with the candidate’s hiring to provide touch points documenting accomplishments and establishing context for activities, especially when including an interdisciplinary focus in Data Science. The communication goes in both directions with the candidate learning the local processes and the Department Chair learning the value and context of the candidate’s accomplishments. I also suggest ongoing discussions between the candidate and fellow faculty members, colleagues, and senior administrators. These provide informal, but important, milestones for progress toward promotion and allow the candidate to build a coherent collection of documentation of accomplishments, context for these accomplishments, and a clear trajectory forward in their career.

In closing, the fields of Statistics/Biostatistics and Data Science are both evolving and dynamic and an academic environment should encourage their growth and development. Academia is an interesting environment with a clear tension between seeking new knowledge but, at times, holding on to established disciplinary distinctions a bit longer than necessary. Managing a healthy and beneficial tension between novelty and establishment requires creativity, patience, collaboration, and experimentation, especially in fostering interdisciplinary excellence in both the current and the next generation of scholars in our own field.

## ACKNOWLEDGEMENTS

Many thanks to Jeff Leek who presented an overview of Data Science faculty activities to a workshop of Department Chairs of Departments of Statistics and Biostatistics at the 2014 Joint Statistical Meetings. During discussions, I suggested that it was important for a Department Chair to package accomplishments in Data Science within a traditional promotion dossier. Jeff subsequently invited me to elaborate on this thought and provide a Department Chair’s perspective in an invited session entitled “The Statistical Identity Crisis: Are we all Data Scientists?” at the 2015 Joint Statistical Meetings. I learned a great deal in preparing the presentation, subsequent reading, and discussions with the other speakers and audience members for that session. Participation in the session encouraged me to think more deeply about the process, ask questions, and attempt to generalize what is often an exercise devoted to one individual at a time.

The opinions expressed above come from my own experiences as a faculty member in three different departments of Biostatistics, and as the chair of one, all within the academic environment of accredited Schools of Public Health, as well as my experiences as a member of two different Promotions and Tenure review committees, as a participant in workshops for chairs of departments of Statistics and Biostatistics sponsored by the American Statistical Association and the Eastern North American Region of the International Biometric Society, as an external reviewer for many individual faculty promotions, and as an external reviewer for departments of both Statistics and Biostatistics. I am grateful for the thoughts and opinions expressed in conversation, emails, presentations, and blog posts by others on related topics. These surely have influenced my own thoughts, and I cite sources when available. While I attempt to speak generally and constructively to bring together these ideas, I accept the responsibility that the opinions expressed below are ultimately my own and am happy to discuss with others as the field continues to develop.

